# The time of strong actomyosin binding depends on electrostatic interactions within the force generating region in human cardiac myosin

**DOI:** 10.1101/2020.04.21.054403

**Authors:** Akhil Gargey, Shiril Bhardwaj Iragavarapu, Alexander V. Grdzelishvili, Yuri E. Nesmelov

## Abstract

Two single mutations, R694N and E45Q, were introduced in the beta isoform of human cardiac myosin to remove permanent salt bridges E45:R694 and E98:R694 in the force-generating region of myosin head. Beta isoform-specific bridges E45:R694 and E98:R694 were discovered in the molecular dynamics simulations of the alpha and beta myosin isoforms. Alpha and beta isoforms exhibit different kinetics, ADP dissociates slower from actomyosin containing beta myosin isoform, therefore, beta myosin stays strongly bound to actin longer. We hypothesize that the electrostatic interactions in the force-generating region modulate affinity of ADP to actomyosin, and therefore, the time of the strong actomyosin binding. Wild type and the mutants of the myosin head construct (1-843 amino acid residues) were expressed in differentiated C_2_C_12_ cells, and duration of the strongly bound state of actomyosin was characterized using transient kinetics spectrophotometry. All myosin constructs exhibited a fast rate of ATP binding to actomyosin and a slow rate of ADP dissociation, showing that ADP release limits the time of the strongly bound state of actomyosin. Mutant R694N showed faster rate of ADP release from actomyosin, compared to the wild type and the E45Q mutant, thus confirming that electrostatic interactions within the force-generating region of human cardiac myosin regulate ADP release and the duration of the strongly bound state of actomyosin.

## Introduction

Myosin II is a nanoscale motor that transduces the chemical energy of ATP to generate force during muscle contraction and perform transport functions in non-muscle cells. ATP binds actomyosin and initiates its dissociation, and ATP hydrolysis drives myosin conformational change, or the recovery stroke, that primes myosin for subsequent strong binding to actin to produce the power stroke. The rate of ATP binding to actomyosin and the rate of ADP release determine duration of the strong actomyosin binding, which is directly related to the average force produced by muscle (Harris and Warshaw 1993). General organization of the motor domain is conserved among myosins (Houdusse and Sweeney 2001), the myosin fold provides a structural frame for the molecular motor function. Myosins have different functions, and their kinetic properties vary dramatically. Kinetics of the actomyosin cycle depend on myosin sequence, and therefore, on inter-residue interactions, which vary in different myosins.

Interatomic forces of physical origin, electrostatics and dispersion, maintain functional interactions between myosin structural elements. Since the dispersion force is a short-range force (r^−6^ dependence), and the electrostatic force between charged residues is the long-range force (r^−2^ dependence), we hypothesize that electrostatic interaction between charged residues of myosin structural elements within the myosin head is the major determinant of myosin kinetics.

In this work we focus on the force-generating region, the region of the myosin head, responsible for significant conformational change during the myosin ATPase cycle. Two helices of the force generation region, the relay helix and the SH1 helix, connect the converter domain with the rest of the myosin head (Figure 1). The relay helix extends from the myosin active site to the converter domain. During the power stroke, the relay helix bends and rotates the converter domain 60° around the SH1 helix (Koppole, Smith et al. 2007).

**Figure 1.**
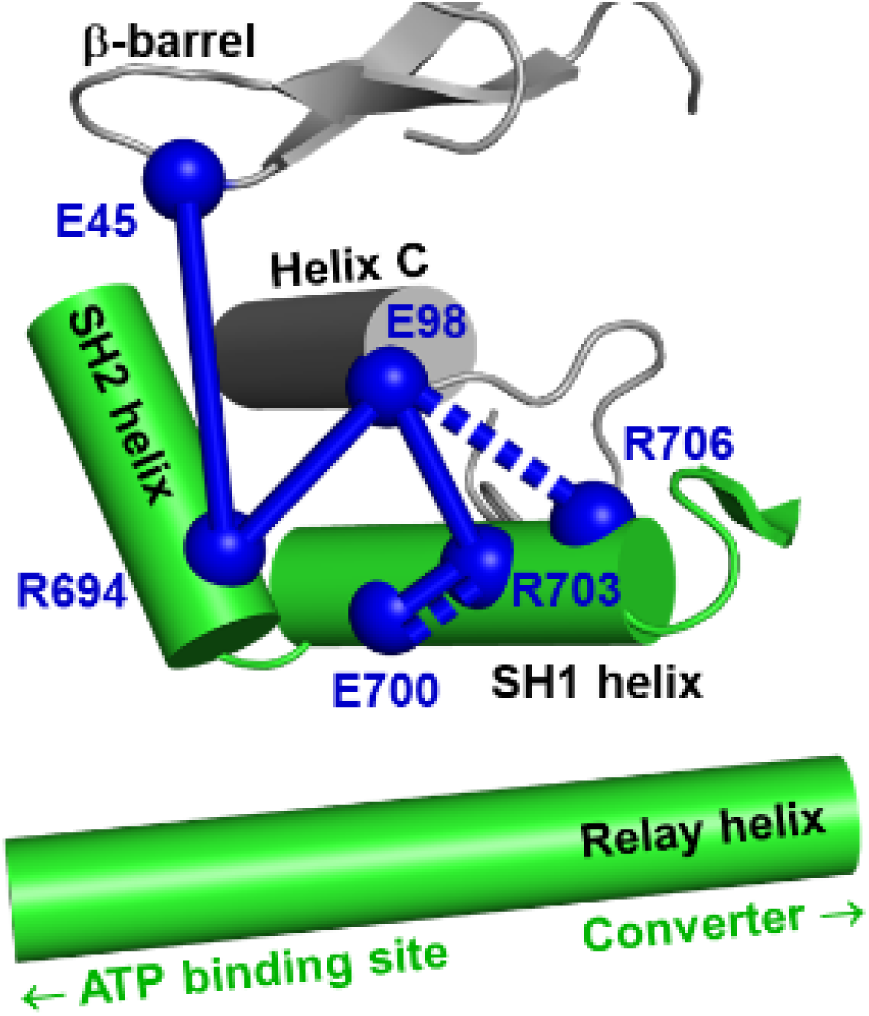
Force-generating region of myosin head. Spheres – charged residues, dashed sticks – permanent α isoform-specific salt bridges, solid sticks – permanent β isoform-specific bridges.

Alpha and beta isoforms of human cardiac myosin share 93% sequence identity of the head domain, but are different kinetically. The alpha isoform (MYH6) is fast and beta isoform (MYH7) is slow. Usually the sequence alignment is used to pick a site for a mutation to examine if the mutation site affects the molecule’s kinetic properties. We chose a different approach. Previously, based on the available crystal structure of the beta isoform (4DB1.pdb, UniProtKB P12883), we built a model of the alpha isoform and ran a molecular dynamics simulation for the 1-843 amino acid residues long constructs of both isoforms (Gargey, Ge et al. 2019). We analyzed obtained trajectories in terms of electrostatic interactions within myosin head. We found significant differences in distribution of electrostatic interactions within the force-generating region of the alpha and beta isoforms (Figure 1). In the beta isoform we found a well-structured electrostatic network of permanent salt bridges, staying put during the whole trajectory. In the alpha isoform, our simulation showed a smaller number of permanent salt bridges, and therefore, higher flexibility of the construct.

The force-generating region is well conserved among myosins and 100% conserved in the alpha and beta isoforms of human cardiac myosin. There are five charged residues in the SH2 and SH1 helices, R694, E700, R703, R706, K707 (MYH7 sequence). Four of these residues form permanent salt bridges in the beta isoform (Figure 1). The effect of mutagenesis of the residues E700, R703, R706 was studied in different myosins before. R706H is the known deafness mutation in nonmuscle myosin IIA (MYH9) (Lalwani, Goldstein et al. 2000), myosin with this mutation shows motility impairment (Iwai, Hanamoto et al. 2006), but undisturbed ATPase activity, indicating that this mutation uncouples the ATPase activity and force production. R703C results in the decreased ATPase activity and reduced actin mobility in the in vitro motility assay (Hu, Wang et al. 2002). E700K mutation (Martinsson, Oldfors et al. 2000) results in slow myosin ATPase activity and slow ATP-induced actomyosin dissociation (Zeng, Conibear et al. 2004). According to our computer simulations, E700 and R703 form a permanent salt bridge, apparently essential for SH1 helix structural integrity during myosin ATPase cycle. We observed this salt bridge in both isoforms. R703 and R706 form a permanent salt bridge with E98 in the beta and alpha isoforms accordingly. Highly conserved E98 is located at the N terminus of the helix C (Cope, Whisstock et al. 1996), and the mutation of the next residue, H97K (MYH7 sequence), introducing positive charge near the negatively charged E98, stops myosin ATPase activity and abolishes actin motility in the in vitro motility assay (Hu, Wang et al. 2002). Myosin mutant H97K cannot bind actin strongly, presumably due to the disturbed release of the products of hydrolysis. Appearance of the positively charged residue next to the negative charge of E98, involved in the electrostatic network, can affect the network, and we suspect that this is the case for the H97K mutant. One can conclude that the electrostatics of the SH1 helix is important for proper myosin functioning, since the SH1 helix is the pivoting helix in myosin power stroke conformational change. It is interesting to mention that disturbed electrostatics not only uncouples myosin ATPase activity and force production (R706), but affects myosin interaction with nucleotide (E700, R703).

Newly developed myosin activator Omecamtiv Mecarbil binds myosin at the SH1 helix (Winkelmann, Forgacs et al. 2015). According to our computer simulations, R694 forms two permanent salt bridges with E98 and E45 in the beta isoform, these salt bridges are not present in the alpha isoform. We hypothesized that removal of one salt bridge (using E45Q mutant) or both salt bridges (using R694N mutant) can modulate kinetics of the beta isoform and in particular the rate of ADP dissociation from actomyosin. We prepared these mutants and studied the effect of R694N and E45Q mutations in the beta isoform of human cardiac myosin to answer the question if electrostatic interactions regulate myosin kinetics, especially the kinetics of the strongly bound state of actomyosin.

## Materials and Methods

### Reagents

*N*-(1-pyrene)iodoacetamide (pyrene) was from Life Technologies Corporation (Grand Island, NY), phalloidin, ATP, and ADP were from Sigma-Aldrich (Milwaukee, WI). All other chemicals were from ThermoFisher Scientific (Waltham, MA) and VWR (Radnor, PA).

### Protein preparation

The β isoform construct of human cardiac myosin motor domain contains 1-843 residues and FLAG affinity tag at the C-terminus. Adenoviruses encoded with the wild type and R694N myosin mutant were purchased from Vector Biolabs (Malvern, PA), amplified using HEK293 cells (ATCC CRL-1573), and purified using CsCl gradient centrifugation. Recombinant human cardiac myosin was expressed in C_2_C_12_ (ATCC CRL-1722) mouse myoblast cells. C_2_C_12_ cells were grown to a 95% confluence on 15 cm diameter plates and infected with optimum dosage of virus determined by a viral-titration assay. Cells were allowed to differentiate post-infection and collected seven days post-infection to extract and purify myosin. Collected cells were washed and lysed in the presence of millimolar concentration of ATP. Cell lysate was incubated with anti-FLAG magnetic beads (Sigma-Aldrich, Milwaukee, WI). Beads were washed and myosin was eluted from the beads by 3x FLAG peptide (ApexBio, Houston, TX). Myosin purity was assessed by Coomassie-stained SDS-polyacrylamide gels and protein concentration was determined by measuring the absorbance at 280 nm using extinction coefficient ε_280nm_ = 93,170 M^-1^cm^-1^, calculated using ProtParam tool of ExPASy web server.

Actin was prepared from rabbit leg and back muscles (Strzelecka-Golaszewska, Prochniewicz et al. 1980, Margossian and Lowey 1982, Waller, Ouyang et al. 1995). F-actin was labeled with pyrene iodoacetamide (Life Technologies Corporation, Grand Island, NY) with the molar ratio 6:1, label:actin. After labeling, actin was cleaned from the excess of label, re-polymerized, stabilized with phalloidin at the molar ratio of 1:1, and dialyzed for two days at T=4°C against the experimental buffer. Concentration of unlabeled G-actin was determined spectrophotometrically assuming the extinction coefficient ε_290nm_ = 0.63 (mg/ml)^−1^cm^-1^ (Houk and Ue 1974). Concentration of labeled G-actin and labeling efficiency were determined spectroscopically using the following expressions: [G-actin]=(A_290nm_ – (A_344nm_·0.127))/26,600 M^-1^ and [pyrene]=A_344nm_ /22,000 M^-1^ (Takagi, Yang et al. 2008). The experimental buffer contained 20 mM MOPS (3-[N-morpholino]propanesulfonic acid) pH 7.3, 50 mM KCl, 3mM MgCl_2_ total concentration. Since log_10_K_A_ for MgATP is 4.29 (Sigel and Song 1996), where K_A_ is the association constant, 3mM MgCl_2_ chelated all ATP used in our experiments, since used ATP concentration was 0.9 mM or less. We do not expect any measurable effect from the KATP complex, since the association constant for such a complex is three orders of magnitude smaller than the constant for MgATP (Melchior 1954). All reported concentrations are final concentrations.

### Acquisition of fluorescent transients

In the ATP-induced actomyosin dissociation experiment, usually 0.5 µM actomyosin was rapidly mixed with ATP solution of variable concentrations. In the competitive inhibition of the ATP-induced actomyosin dissociation experiment, 0.5 µM actomyosin was rapidly mixed with the premixed ATP and ADP solution. The concentration of ATP in solution was 0.6 mM or 0.9 mM and the concentration of ADP varied from 20 µM to 200 µM. Transient fluorescence of pyrene-labeled actin was measured with a Bio-Logic SFM-300 stopped flow transient fluorimeter (Bio-Logic Science Instruments SAS, Claix, France), equipped with a FC-15 cuvette. The pyrene fluorescence was excited at 365 nm and detected using a 420 nm cutoff filter. Multiple transients were acquired and averaged to improve signal to noise ratio. 8000 points were acquired in each experiment. All experiments were performed at T=20° C.

### Analysis of fluorescence transients

The transients obtained in each experiment were fitted by the single exponential function S(t) = S_o_+A·exp(-k_obs_·(t-t_0_)), or the double exponential function S(t) = S_o_+A_1_·exp(-k_obs1_·(t-t_0_)) + A_2_·exp(-k_obs2_·(t-t_0_)). S(t) is the observed signal at the time t, A_i_ is the signal amplitude, t_0_ is the time before the flow stops, and k_obsi_ is the observed rate constant. Transients, obtained for the same actomyosin preparation at different concentrations of the nucleotide were fitted together, assuming the known value of t_0_, measured in a separate experiment, and the constant value of S_0_, which depends on the concentration and labeling efficiency of pyrene-labeled actin in the actomyosin preparation. In the case of the two-exponential global fit, we kept amplitudes of the transients constrained, A_1_+A_2_=const, to account for the conservation of mass. Transients of the ATP induced actomyosin dissociation were fitted with the one-exponential function to determine the rate constant k_obs_. The dependence of the observed rates on the ATP concentration was fitted by a hyperbola, v = V_max_·[ATP]/(K_app_ +[ATP]), allowing the determination of the maximum rate, V_max_ (the horizontal asymptote). In the case of the ATP-induced actomyosin dissociation, the rate constant k_+2T_ is the V_max_ and the equilibrium constant of the collision complex formation K_1T_ is 1/K_app_. To determine the bimolecular rate (K_1T_k_+2T_), the dependence of the observed rates on the ATP concentration was fitted by a straight line at small concentrations of ATP. Transients of the competitive inhibition experiment were fitted with the two-exponential function, where the rate constant of the fast process reflects the ATP-induced actomyosin dissociation, and the slower process is the inhibition of the ATP-induced actomyosin dissociation by ADP, reporting the rate of ADP release from actomyosin. Following (De La Cruz and Ostap 2009), the observed rate of the slow exponential component k_obs_ depends on the rate of ADP dissociation, k_-1D_, normalized on the probability of ATP binding to actomyosin

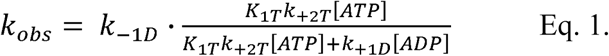

where the bimolecular rate K_1T_k_+2T_ is determined in the experiment without ADP. The fit allows the rate constants of ADP binding to and dissociation from actomyosin, k_+1D_ and k_-1D_, to be determined. These rate constants were measured at the close to saturated ADP and ATP concentrations, the transients were fitted to the two-exponential function, with the rate constant of the fast exponent fixed at the rate of actomyosin dissociation, to account for the non-complete saturation with ADP. Rate constants were obtained in experiments with proteins from at least three independent preparations. Obtained rate constants were averaged, and mean values and standard errors were used to fit with Equation 1. All data fits were performed with Origin 8 (OriginLab Corp, Northampton MA). The statistical significance of results was tested with ANOVA integrated in Origin 8 software. A significance level of P < 0.05 was used for all analyses.

## Results

### Salt bridges in the force-generating region of myosin S1

In our previous computer simulations (Gargey, Ge et al. 2019) we identified several permanent salt bridges in the force-generating region of human cardiac myosin, beta isoform (Figure 1), which form isoform-specific electrostatic network. The bridge E700-R703 exists in both isoforms, the bridge E98-R703 in beta isoform switches to the bridge E98-R706 in alpha isoform, and two bridges, E98-R694 and E45-R694, exist only in the beta isoform. Previous studies confirmed the importance of charged residues in this region (Lalwani, Goldstein et al. 2000, Martinsson, Oldfors et al. 2000, Hu, Wang et al. 2002, Zeng, Conibear et al. 2004, Iwai, Hanamoto et al. 2006). According to our computer simulations, all these residues are involved in the permanent electrostatic network in the force-generating region. The bridge E98-R694 links the SH2 helix and the N terminus of the helix C. The bridge E45-R694 links the SH2 helix and the beta barrel of the N-terminus of myosin head.

### Design and preparation of myosin S1 constructs

We choose residues E45 and R694 for mutagenesis because of their participation in the isoform-specific electrostatic interactions within the force-generating region of the myosin head. The roles of other charged residues in the region were studied previously (Lalwani, Goldstein et al. 2000, Martinsson, Oldfors et al. 2000, Hu, Wang et al. 2002, Zeng, Conibear et al. 2004, Iwai, Hanamoto et al. 2006). We prepared constructs of the wild type and two mutants, E45Q and R694N, of the human cardiac myosin motor domain (1-843 amino acid residues long) in the beta isoform background. All constructs have the FLAG tag at the C-terminus. We used differentiated C_2_C_12_ cells to express constructs and FLAG tag affinity chromatography to purify expressed constructs. Purity of the expressed constructs was confirmed with SDS PAGE (Figure 2).

**Figure 2.**
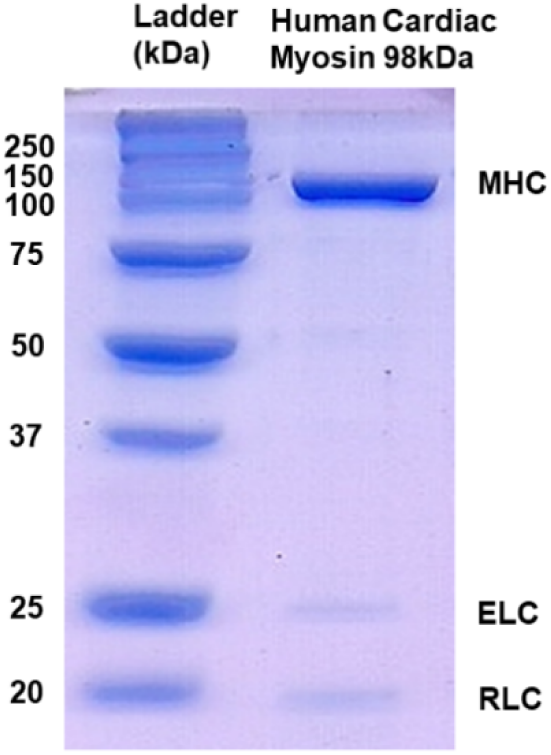
SDS-PAGE of purified recombinant myosin head. 98 kDa human cardiac S1 co-purifies with murine ELC and RLC.

### ATP-induced actomyosin dissociation

We monitor ATP-induced actomyosin dissociation using fluorescence of pyrene labeled actin. In the absence of ATP, actin strongly binds myosin and pyrene fluorescence is quenched. Pyrene fluorescence increases upon actomyosin dissociation and formation of the weakly bound actomyosin complex (Kouyama and Mihashi 1981, Criddle, Geeves et al. 1985). Therefore, pyrene fluorescence reports the lifetime of the strongly bound state of actomyosin. ATP binds actomyosin in a two-step process (Scheme 1). The first step is the rapid equilibrium, when strongly bound actomyosin forms a collision complex with ATP, and k_-1T_ >> k_+2T_. Pyrene fluorescence does not change during this step. Upon isomerization of the actomyosin·ATP collision complex, ATP binding results in irreversible dissociation of actomyosin. In our experiments, formation of the actomyosin complex (prepared by mixing of expressed myosin constructs with the pyrene-labeled rabbit skeletal actin) led to decreased pyrene fluorescence. This decrease was similar for all myosin constructs, indicating strong binding of actin and expressed myosin constructs. When prepared actomyosin is rapidly mixed with ATP, the time course of pyrene fluorescence follows single exponential kinetics. Figure 3 shows the fluorescence transient observed for the WT myosin at 20°C when 0.5 µM actomyosin is mixed with 900 µM ATP in the stopped-flow fluorimeter (all reported concentrations are in the final mixture, here and throughout the text). The rate is 449.7 s^-1^, in excellent agreement with the published data (Deacon, Bloemink et al. 2012). The observed rate constants for the WT myosin and the mutants depend on ATP concentration hyperbolically (Figure 4). The horizontal asymptote of the hyperbolic fit of the kinetic rates gives the maximum velocity of the ATP-induced actomyosin dissociation (491.5±42.8 s^-1^, 338.8±27.6 s^-1^, and 618.5±43.5 s^-1^ for the WT, R694N and E45Q mutants accordingly). The dissociation constants of the collision complex, determined from the fit, were 220.3±38.5 µM, 127.4±24.0 µM, and 137.4±21.7 µM. The second-order association rate constant (K_1T_k_+2T_) is determined at small concentrations of ATP (Figure 5), when dependence of the reaction rate on ATP is linear (Segel 1976). The association constants are 2.11±0.12 µM^-1^s^-1^, 1.85±0.02 µM^-1^s^-1^, and 3.27±0.18 µM^-1^s^-1^ for the WT, R694N and E45Q mutants. The final amplitudes of all transients, obtained for the same preparation of actomyosin, were the same for all used concentrations of ATP, showing complete dissociation of actomyosin complex and confirming that the ATP-induced dissociation is irreversible.

**Scheme 1.**
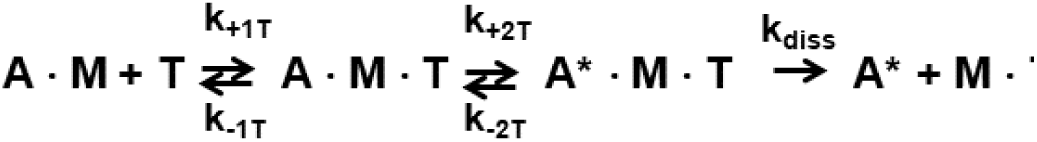
ATP-induced actomyosin dissociation. A = pyrene labeled actin, M = myosin, T = ATP. A* = actin with unquenched pyrene fluorescence. k_diss_ >> k_+2T_ + k_-2T_.

**Figure 3.**
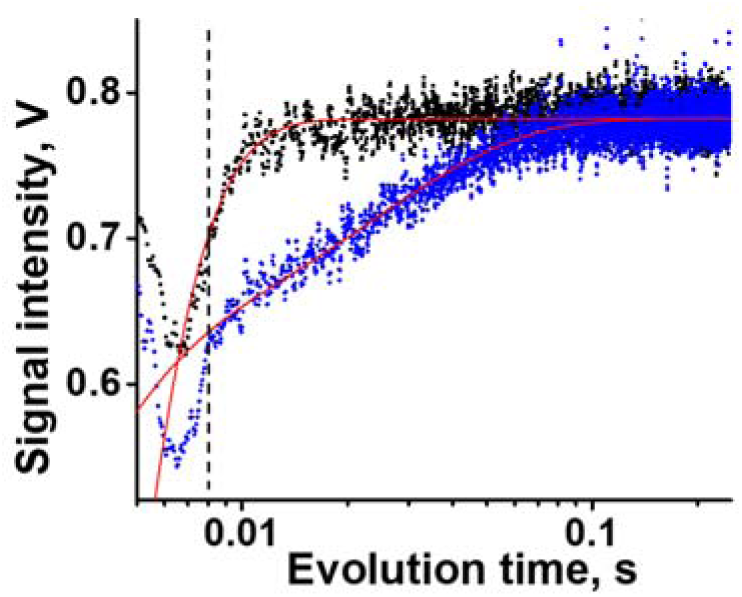
Typical transients in the experiment of the ATP-induced actomyosin dissociation with and without ADP. Actomyosin (0.5 µM) rapidly mixed with ATP (upper trace), or the mixture of ATP and ADP (lower trace). [ATP] = 900 µM in both cases. [ADP] = 200 µM when present in the mixture. All transients for the same protein preparation are fitted globally to the two-exponential equation, except for the [ADP]=0 transient, fitted to the one-exponential equation. Dead time, measured in a separate experiment, constrains the fit (all fits intercept at the mixing time in the bottom left corner). Vertical dashed line shows the time of the flow stop and the beginning of the fit. Meaningful kinetic traces lay on the right side of the dashed line. On the left there are flow artefacts.

**Figure 4.**
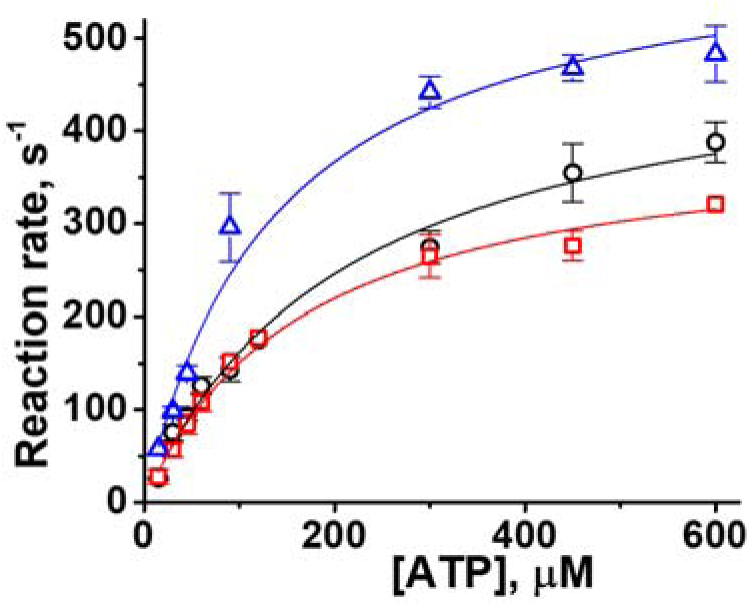
Rate of ATP-induced actomyosin dissociation. Circles, WT, N=3, squares, R694N mutant, N=3 (the point at [ATP]=600 μM, N=1), triangles, E45Q mutant, N=3. Reaction rates fitted with a hyperbola, k_+2T_ = 491.5±42.8 s^-1^, 338.8±27.6 s^-1^, 618.5±43.5 s^-1^ for WT, R694N, and E45Q respectively.

**Figure 5.**
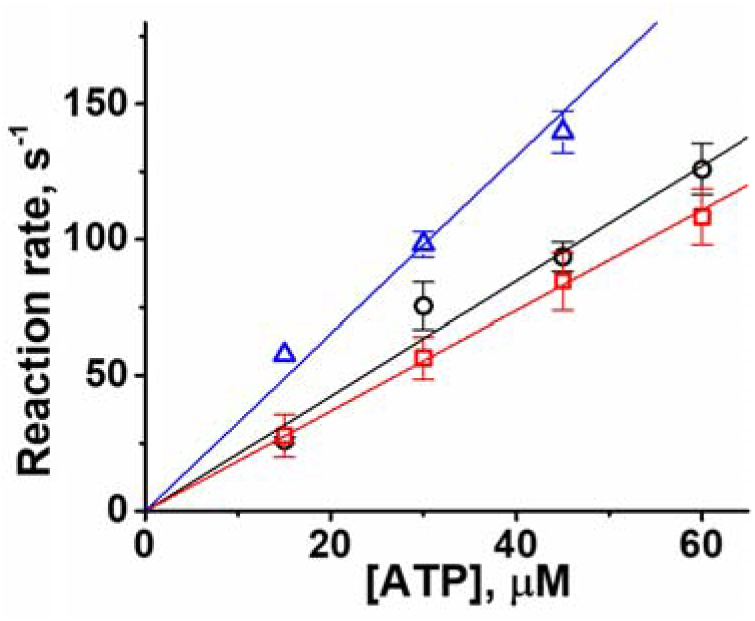
ATP-induced actomyosin dissociation. Observed reaction rates at low [ATP] are fitted by a straight line, the second order reaction rate constant is determined from the slope of the line. Circles, WT, N=3, squares, R694N mutant, N=3, triangles, E45Q mutant, N=3. K_1T_k_+2T_ = 2.12±0.12 M^-^ 1s^-1^, 1.85±0.02 M^-1^s^-1^, 3.27±0.18 M^-1^s^-1^ for WT, R694N, and E45Q respectively.

### ADP dissociation from actomyosin

ADP has high affinity to the WT human cardiac actomyosin. The equilibrium dissociation constant of ADP is in the micromolar range (Deacon, Bloemink et al. 2012, Gargey, Ge et al. 2019). To measure the rate of ADP dissociation from actomyosin we rapidly mix pyrene-labeled actomyosin with premixed ADP and ATP. In our experiments we kept ATP concentration constant (near saturation, but not saturated, 600 µM or 900 µM) and vary ADP concentration from 20 µM to 200 µM. Upon mixing, actomyosin can bind either ATP or ADP and form either an actomyosin·ATP or actomyosin·ADP complex (Scheme 2). ATP binding results in actomyosin dissociation and in increased pyrene fluorescence. ADP binding does not dissociate actomyosin and therefore does not change the intensity of pyrene fluorescence. If actomyosin, actomyosin·ATP, and actomyosin·ADP complexes are in rapid equilibrium, we should observe one-exponential increase of pyrene fluorescence, corresponding to the competitive inhibition of the ATP-induced actomyosin dissociation. At saturated concentration of ATP there should be no ADP dependence on the rate of actomyosin dissociation (Segel 1976). In our experiments with the constructs of human cardiac myosin we usually observe two-exponential kinetics of ATP-induced actomyosin dissociation in the presence of ADP (Figure 3). Our observation of the two-exponential kinetics suggests that there is no rapid equilibrium in the mixture of actomyosin, ATP, and ADP. ADP readily binds actomyosin, and the rate of ADP dissociation from actomyosin is slower than the rate of ATP binding. The amplitude of the fast component decreases with the increase of ADP concentration in the mixture. This decrease of the amplitude reflects decrease in the probability that actomyosin binds ATP. The rate of the fast component of the observed two-exponential kinetics does not depend on ADP concentration and corresponds well to the rate of ATP-induced actomyosin dissociation. We conclude that there is no fast exchange between ADP and the actomyosin·ADP complex in the beta isoform actomyosin, wild type, and mutants. The slow component of the two-exponential transient depends on the concentration of ADP and reflects ADP binding to actomyosin, followed by ADP dissociation and subsequent ATP binding, resulting in actomyosin dissociation. The rate of ADP dissociation from actomyosin can be obtained from the fit of the experimentally observed rates of the slow component to the Equation 1, using measured ATP binding constant K_1T_k_+2T_, and known concentrations of ATP and ADP (Figure 6). The fit gives the rate k_-1D_, as well as the rate of ADP binding to actomyosin, k_+1D_. We found that the rate of ADP dissociation from actomyosin is the same within the error of experiment for the WT and E45Q constructs, and 65% faster for the R694N myosin construct compared to the WT.

**Scheme 2.**
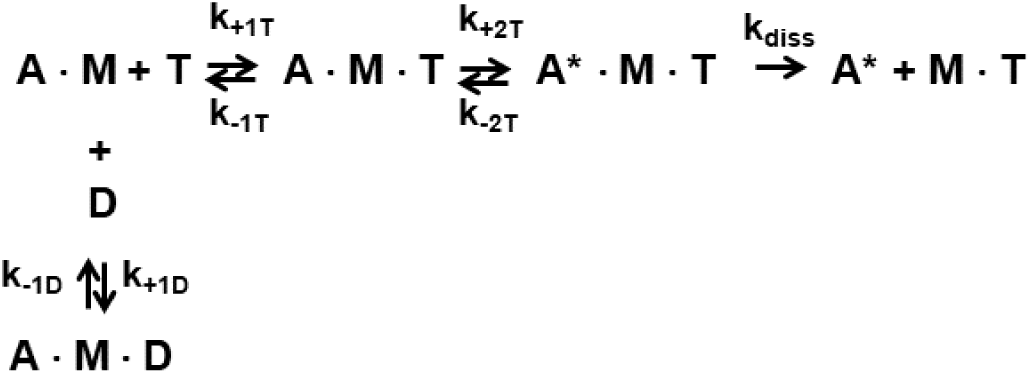
ATP-induced actomyosin dissociation, competitive inhibition with ADP.

**Figure 6.**
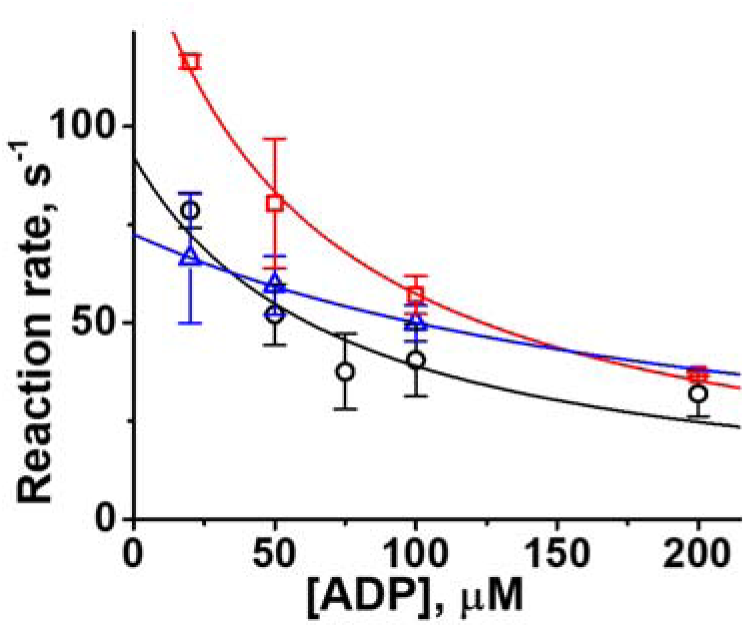
ATP induced ADP dissociation from actomyosin. Circles, WT, squares, R694N mutant, [ATP] = 900 μM, triangles, E45Q mutant, [ATP] = 600 μM, N = 3. Lines, fit to the Eq. 1. The rate of ADP dissociation from actomyosin is k_-1D_ = 92.3±13.2 s^-1^, 152.7±7.7 s^-1^, and 76.4±6.0 s^-1^ for the WT, R694N, and E45Q constructs respectively.

## Discussion

In this study we experimentally verified our hypothesis that electrostatic interactions in the force-generating region of the myosin head modulate the rate of ADP dissociation from actomyosin. A single mutation, R694N, disrupts the salt bridge, R694:E98, which exists only in the slow beta isoform of human cardiac myosin. The mutant R694N shows faster rate of ADP dissociation from actomyosin, and therefore, shorter duration of the strongly bound actomyosin state, t_s_. The timing of the strongly bound state depends on how fast actomyosin enters and exits the state, t_s_ = 1/k_+2T_ + 1/k_-1D_, where k_+2T_ is the rate of ATP-induced actomyosin dissociation, and k_-1D_ is the rate of ADP dissociation from actomyosin. For the WT beta isoform human cardiac myosin t_s_ = 12.9±1.6 ms, for the R694N mutant t_s_ = 9.5±0.4 ms, 26% shorter. Another isoform-specific salt bridge R694:E45, examined in this work, apparently does not play any major role in the regulation of the strongly bound state of actomyosin, t_s_ = 14.7±1.0 ms for the E45Q mutant. Structural details of such regulation of myosin kinetics are yet to be determined.

ADP binds actomyosin faster than ATP (Table 1), apparently reflecting the difference in charge of these molecules. At pH 7 and higher, both MgADP and MgATP are ionized (Phillips, George et al. 1966), and the total charge of MgADP and MgATP is (-1e), and (-2e) accordingly. The charge dependence of the kinetics of nucleotide binding suggests an overall negative charge of the myosin active site, and therefore electrostatic repulsion when nucleotide binds actomyosin.

**Table 1.** Actomyosin kinetic rate constants, mean±SE. The data are averages of three independent protein preparations, T=20°C.

|  | WT | R694N | E45Q |
| --- | --- | --- | --- |
| Actomyosin dissociation |  |  |  |
| $K_{1T}k_{+2T}, \mu\text{M}^{-1}\text{s}^{-1}$ | 2.12±0.12 | 1.85±0.02 | 3.26±0.18 |
| $k_{+2T}, \text{s}^{-1}$ | 491.5±42.8 | 338.8±27.6 | 618.5±43.5 |
| Actomyosin collision complex formation |  |  |  |
| $K_{1T}, \text{mM}^{-1}$ | 4.5±0.8 | 7.8±1.5 | 7.3±1.1 |
| ADP binding to and release from actomyosin |  |  |  |
| $k_{+1D}, \mu\text{M}^{-1}\text{s}^{-1}$ | 26.0±8.2 | 27.6±2.9 | 10.5±3.6 |
| $k_{-1D}, \text{s}^{-1}$ | 92.3±13.2 | 152.7±7.7 | 76.4±6.0 |
| ADP dissociation from actomyosin |  |  |  |
| $K_{1D}, \mu\text{M}$ | 3.5±1.2 | 5.5±0.6 | 7.3±2.6 |

The diffusion-controlled rate of ATP and actomyosin association is k_+1T_ = 255µM^-1^s^-1^, this rate is calculated using the expression k = 4·D_ATP_·*a* (Shoup, Lipari et al. 1981), where *a* is the radius of myosin active site (we assume it equals to 0.4 nm, the radius of ATP), and D_ATP_ is the diffusion coefficient of ATP, D_ATP_ = (2.65±0.09)·10^−10^ m^2^s^-1^ at T=20°C (Ge, Gargey et al. 2019). Assumption of the larger diameter of the active site makes the rate of association faster, therefore, the value k_+1T_ = 255 µM^-1^s^-1^ is the minimal value for the rate of the diffusion-controlled reaction of ATP and actomyosin. The expression k = 4·D_ATP_·*a* follows from the consideration of the diffusion-controlled bimolecular reaction between the large macromolecule with a single ligand-binding site and a small uniformly reactive ligand (Solc and Stockmayer 1971, Shoup, Lipari et al. 1981). The expression is accurate when (a) the diffusion of actomyosin is much slower that the diffusion of ATP, (b) the active site is a small part of the surface of actomyosin, and (c) the diameter of ATP is much smaller than the diameter of the actomyosin complex.

The hyperbolic dependence of ATP-induced actomyosin dissociation indicates that ATP binds actomyosin in two steps, first forming the collisional actomyosin·ATP complex, which is in rapid equilibrium with ATP and actomyosin, and then binding actomyosin practically irreversibly, causing actomyosin dissociation. Both mutants and the WT myosin construct exhibit that hyperbolic dependence. The value of the equilibrium constant of the collision complex formation K_1T_ is two times larger for the mutants, compared to the WT, indicating higher affinity of ATP to actomyosin in the mutants. Assuming millimolar concentration of ATP in muscle (Kushmerick, Moerland et al. 1992), increase of the equilibrium constant from 4 mM^-1^ to 7 mM^-1^ virtually does not change the population of the actomyosin·ATP complex. For the WT myosin, assuming the diffusion-controlled rate of the collision complex formation (k_+1T_ = 255uM^-1^s^-1^), and experimentally measured equilibrium constant K_1T_ = 4 mM^-1^, the rate of the reverse reaction is k_-1T_ = k_+1T_/K_1T_ = 64·10^3^ s^-1^. The same analysis gives k_-1T_ = 36·10^3^ s^-1^ for both mutants. These correspond well to the rapid equilibrium condition of the reaction of ATP-induced actomyosin dissociation, taking into account that the rate of the next step (step 2 in the Scheme 1), determined in this study, is 350 – 600 s^-1^ (Table 1). The half-lifetime of the collision complex is then t_1/2_ = ln(2)/k_-1T_ = 11 µs for the WT myosin and 19 µs for both mutants.

If the active site is not open all the time and is in equilibrium between the open and closed states, the rate of the collision complex formation k_+1T_ can be smaller than the rate of the diffusion-controlled reaction. Then, the reverse rate k_-1T_ decreases according to the equilibrium constant K_1T_, determined in the experiment. The reverse rate k_-1T_ should be faster than the rate k_+2T_ to satisfy to the observed hyperbolic dependence of ATP-induced actomyosin dissociation. For WT myosin, if k_+1T_ = 25.5 uM^-1^s^-1^, an order of magnitude smaller than the rate of the diffusion-controlled reaction, k_-1T_ = 6.4·10^3^ s^-1^, that still corresponds to the rapid equilibrium condition of the step 1 in the Scheme 1. If k_+1T_ = 2.5 uM^-1^s^-1^, the rapid equilibrium condition is not satisfied since the reverse rate k_-1T_ becomes of the same order of magnitude as the forward rate k_+2T_. Since the equilibrium constant K_1T_ is higher for mutants, the decrease of the forward rate k_+1T_ results in the even slower reverse rate, k_-1T_. Therefore, to correspond to the experimentally obtained hyperbolic dependence of ATP-induced actomyosin dissociation, the rate of the collision complex formation should be in the range of k_+1T_ = 255 uM^-1^s^-1^ – 25 uM^-1^s^-1^. Then, the reverse rate is in the range of k_-1T_ = 64·10^3^ s^-1^ – 6.4·10^3^ s^-1^ for the WT and k_-1T_ = 36·10^3^ s^-1^ – 3.6·10^3^ s^-1^ for both mutants.

Mutagenesis in the force-generating region affects the rate of actomyosin dissociation k_+2T_. The rate change is in the range of ±30% for WT myosin and mutants. The rate of actomyosin dissociation is faster than the rate of ADP dissociation from actomyosin for all studied myosin constructs. Therefore, the duration of the strongly bound actomyosin state is determined mostly by the rate of ADP dissociation from actomyosin.

When we mix actomyosin with the mixture of ATP and ADP, the actomyosin dissociation follows two-exponential kinetics. This indicates that there is no rapid equilibrium between actomyosin and the actomyosin·ADP complex. The fast exponential component of the reaction corresponds to the ATP-induced actomyosin dissociation, and the slow component corresponds to ADP binding to actomyosin and therefore delayed ATP-induced actomyosin dissociation. There is no [ADP] dependence of the fast component of actomyosin dissociation, this confirms the absence of the rapid equilibrium between actomyosin and the actomyosin·ADP complex. To determine the true rates of ADP binding to and dissociation from actomyosin, we weight observed rates of the slow exponential component with the probability of the subsequent ATP binding, as it was suggested before (Eq. 1, (Robblee, Olivares et al. 2004, Robblee, Cao et al. 2005, De La Cruz and Ostap 2009)). The rates of ADP binding to actomyosin are slower than the diffusion-controlled rate of the reaction (Table 1). The rate of ADP dissociation from actomyosin is the same for the WT myosin and E45Q mutant, and 65% faster for the R694N mutant. Observed rates are close to the rates reported for cardiac myosin S1 from different species (Marston and Taylor 1980, Siemankowski and White 1984, Siemankowski, Wiseman et al. 1985, Palmiter, Tyska et al. 1999), and close to the kinetic difference between alpha and beta isoforms of cardiac myosin (Palmiter, Tyska et al. 1999). The result for myosin mutant R694N supports our hypothesis that electrostatic interactions within the myosin head modulate its kinetics.

Since ADP is similar to ATP in size and structure, we may suggest that ADP binds in the two-step mechanism (Scheme 3), similar to ATP. The hypothesis of the several sequential actomyosin·ADP states in the actomyosin cycle was discussed before (Rosenfeld, Xing et al. 1998, Hannemann, Cao et al. 2005, Bloemink and Geeves 2011). The upper limit of the rate of ADP binding to actomyosin is the rate of the diffusion-controlled reaction, and the lower limit is the rate of the reaction obtained in our experiments (Table 1). The rate of ADP dissociation from actomyosin must be at least an order of magnitude slower than the similar rate for ATP, because of the absence of the rapid equilibrium for actomyosin and actomyosin·ADP complex. Therefore, the upper estimate for the reverse rate is k_-1D_ = 64·10^2^ s^-1^ for the WT and k_-1D_ = 36·10^2^ s^-1^ for both mutants. The lower estimate of the rate is the rate of ADP dissociation from actomyosin, obtained in our experiments (Table 1). Assuming that K_1D_K_2D_ in Scheme 3 is the ratio of the determined rates of complex formation and dissociation, K_1D_K_2D_ = k_+1D_/k_-1D_ = 0.28±0.10 µM^-1^s^-1^ for the WT and 0.18±0.02 µM^-1^s^-1^ and 0.14±0.05 µM^-1^s^-1^ for the R694N and E45Q mutants accordingly. If the rate of ADP binding to actomyosin is diffusion-controlled (k_+1D_ = 255 µM^-1^s^-1^), and the reverse rate is in the range of k_-1D_ = 64·10^2^ s^-1^ – 92 s^-1^, then K_2D_ = K_1D_K_2D_ ·k_-1D_/k_+1D_ = 7.0 – 0.1 for the WT myosin. Discussed above the range of the rate of ADP binding to actomyosin (Scheme 3, k_+1D_ = 255 µM^-1^s^-1^ – 26 µM^-1^s^-1^) and the reverse rate (Scheme 3, k_-1D_ = 64·10^2^ s^-1^ – 92 s^-1^), extends the range of K_2D_ for the WT myosin to 68.9 – 0.1. The same consideration for the R694N and E45Q mutants results in the range for the equilibrium constant K_2D_ in ranges 23.4 – 0.11 and 48.0 – 0.04 accordingly. Decrease of the equilibrium constant K_2D_ for mutants reflects decreased population of the second actomyosin·ADP state, if the two-step kinetics of the actomyosin·ADP complex is considered.

**Scheme 3.**
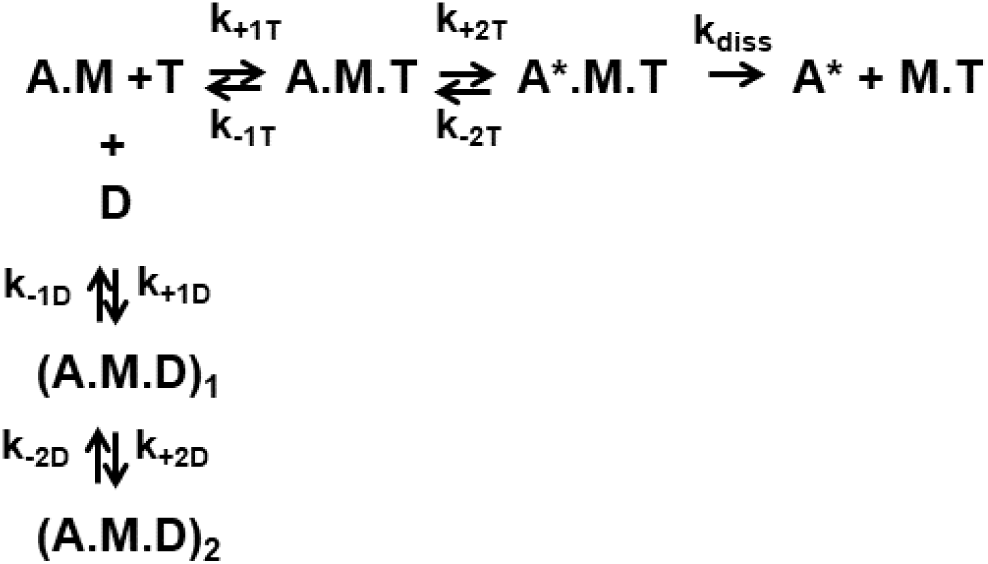
ATP-induced actomyosin dissociation, inhibition with ADP. Two sequential states of the actomyosinADP complex considered.

### Conclusion

In this work we experimentally confirmed that electrostatic interactions within the head of human cardiac myosin modulate the time of the strongly bound state of actomyosin.

## Compliance with ethical standards

Actin was produced from rabbit skeletal tissue. All experimental protocols were approved by the Institutional Animal Care and Use Committee of UNC Charlotte and all experiments were performed in accordance with relevant guidelines and regulations.

## Acknowledgements

This work was supported by National Institutes of Health grant HL132315

